# Biochemical network motifs can transduce and process oscillatory information

**DOI:** 10.1101/2022.12.10.519932

**Authors:** Jamil Reja, Tess K. Fallon, Andrew M. Leader, Evren U. Azeloglu

**Affiliations:** Division of Nephrology, Department of Medicine, Icahn School of Medicine at Mount Sinai; Department of Pharmacological Sciences, Icahn School of Medicine at Mount Sinai, New York, NY; Department of Electrical Engineering, Columbia University, New York, NY

## Abstract

Biological networks that are formed through amalgamation of signaling pathways include recurrent configurations called network motifs. These statistically over-represented subgraphs are often formed through interconnected enzyme-substrate relationships that are known to result in highly dynamic downstream behavior, including oscillatory output. Such signals are abundant in biology: heartbeats, circadian rhythms, and cell cycles all exhibit characteristic frequencies. Though there has been great emphasis on how oscillations can be generated through network dynamics, little is known about oscillatory information processing or transduction capacity of network motifs. We employ ordinary differential equations-based dynamical modeling to understand how different network topologies impact oscillatory signal propagation through a multi-enzyme network. We model enzyme-substrate interactions of 20 commonly observed motifs using Michaelis-Menten kinetics. We then perform deterministic Monte Carlo simulations using a range of biologically relevant enzymatic parameters and input frequencies. From these simulations, we quantify signal propagation characteristics using cluster analysis, categorize different motif responses based on output characteristics, and identify potential mechanisms for oscillatory signal processing using parameter sensitivity analysis. We see that the input-output responses depend on network topology and enzyme kinetic parameters. Enzymatic motifs show median oscillatory suppression of 30-135 decibels, with three-node coherent feedforward loops showing the lowest propensity for oscillatory signal suppression. Motifs that contained negative feedback or four-node coherent feedforward loops had the biggest potential to act as AC-to-DC converters, translating oscillatory input signals into transient impulses or sustained continuous outputs, respectively. We conclude that enzyme networks can process and decode information within oscillatory inputs in a frequency- and network-dependent manner.

## Introduction

Signal transduction is the mechanism through which information is relayed from one courier to another. In cellular biology, couriers of information are often presumed to be proteins or small molecules.^1,2^ In reality, however, the transmitted information can be biochemical, mechanical, or electrical, and such information can be relayed through activity state, concentration, spatial context or activation duration of biological molecules.^3^ In fact, the temporal behavior of a signal is just as important as its strength or spatial localization.^4^ *Dictyostelium discoideum*, for example, determines its motion trajectory based on chemotaxis, which depends on the frequency of cyclic bursts of cAMP that control the nuclear shuttling of GATA.^5^ Similarly, the spatial context of the molecules could substantially modify information flow. For example, activation of integrin receptors through mechanical signals, leading to long-term changes in epithelial cell polarity, depends on spatially cyclic changes in surface area-to-volume ratio.^6^

It has been well established that adapter proteins and enzymes responsible for processing biological signals can be modelled in subgraphs known as network motifs.^7,8^ Two-, three- and four-node network motifs are most commonly studied, as well as most ubiquitously present in nature.^9,10^ Although these simple motifs can form large networks that can carry out highly complex decision-making processes,^11^ even the simplest forms, such as a two-node positive feedback loop, could result in complex phenomena, such as bi-stability.^12,13^ While such versatility may be ideal for generating complex behavior, when combined with contextual modifiers, the same flexibility may also result in biological functions being overtly fragile.^14,15^ It has also been found that most cellular information processing is carried out dynamically, from upstream to downstream, within a signal pathway. After processing, the final output can be highly variable.

Some of the most abundant outputs in biology, such as heartbeats, circadian rhythms, and cell cycles, can be described as oscillatory outputs.^16,17,18^ There has been great emphasis on how these signals can be generated through network motif dynamics; however, little is known about how oscillatory information is processed when it acts as an input for a downstream network motif. It has been shown that incoherent forward loops have the ability to process oscillatory signals.^27^ However, many other network motifs exist in biological systems. In this paper, we aim to answer questions regarding how biological systems process oscillatory information as an input source. Some of the other questions we try to address are: are particular network motifs capable of processing oscillatory inputs; if so, what enzymatic parameters or innate biological characteristics are required to carry out such cellular signal transduction; lastly, do the capabilities of network motifs vary based on the characteristics of the input signal (e.g., input signal frequency and/or amplitude, enzyme properties, etc.)?

Using systems of ordinary differential equations (ODEs), we devised a computational approach to model a range of network motifs and parameters. We then assessed and clustered individual motif responses to varying oscillatory signal inputs in an attempt to understand the dynamic signal processing capabilities of various network motifs. Simulations show that enzymatic networks with biologically reasonable kinetic parameters are indeed capable of processing oscillatory information, leading to various types of non-oscillatory outputs. Our results suggest that biochemical enzymes, depending on their context, could be responsible for complex and biologically meaningful responses to cyclic inputs.

## Methods

Twenty of the most common biochemical network motifs (**S1 Fig**) were modeled computationally using ODEs.^7,19^ These ODEs implemented the well-known Michaelis-Menten model of enzyme kinetics (Eq 1-2).

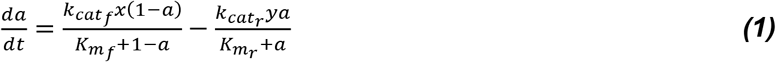

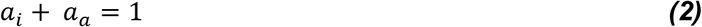

where ***a*** represents the concentration of a substrate (node) within the motif. Subsequently, ***da/dt*** models the change in that molecular substrate with respect to time. ***x*** is the sinusoidal input signal. ***k_cat_*** and *K_M_* are enzyme properties, which indicate the maximum turnover rate of the reaction and the substrate affinity to the enzyme, respectively.

To generate the varying motif structures, either a positive activation relationship (denoted by arrow, ⟹) or a negative inhibitory relationship (denoted by plunger, --|) was randomly generated between each node (**S1 Fig**). For each simulation, a random ***k_cat_*** and *K_M_* were assigned for each substrate-enzyme interaction. ***k_cat_*** values were log-uniform generated on the range [10^-1^, 10^2^]/s/s.^22^ *K_M_* values were also loguniform generated, but on the range [10^-2^, 10^3^] μM. All non-labelled reactions are catalyzed by a constant amount of enzyme with a constant ***k_cat_*** and **K_m_**. Constant enzyme amounts are set to 0.5 μM with constant ***k_cat_*** set to 10^0.5^ per second (approximately 3.16 per second) and constant K_M_ values are set to 10^0.5^ μM. Here, we also assume there is total conservation of enzyme/protein, **a**, and the sum total is 1. Equations for all motifs are outlined in **S2 Appendix**.

Computational generation of data and subsequent analysis were done in MATLAB 2016Ra. A single simulation solves a system of ODEs for a single motif with randomly generated ***k_cat_*** and ***K_m_*** and a single non-variable frequency (**Fig 1A**). For each motif, we first determine the steady state activity levels through an extensive equilibrium period of 100 hours (**t** = 3.6 x 10^5^ seconds) with a normalized input of ***x*** = 0.5*μM*. The initial values of the system are then set to these steady state equilibrium values, after which we prescribe a sinusoidal input stimulus

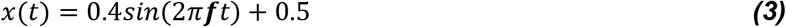

where ***f*** is one of seven biologically relevant frequencies, 0.001 Hz, 0.003 Hz, 0.01 Hz, 0.03 Hz. 0.1 Hz, 0.3 Hz, 1 Hz (**Fig 1B, C, D**). We note that it is necessary to specify time vectors for the simulations. ***t*** is the time vector used to solve the ODE system, and is prescribed as

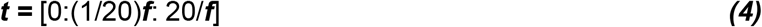

where ***f*** is the frequency of the input stimulus. The core data was generated on the time vector denoted in Equation 4 to optimize file size efficiency. For particularly interesting simulations, where a higher temporal resolution was beneficial, we solved the systems over a time vector, independent of frequency:

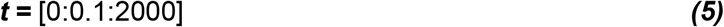

**Fig 1.**
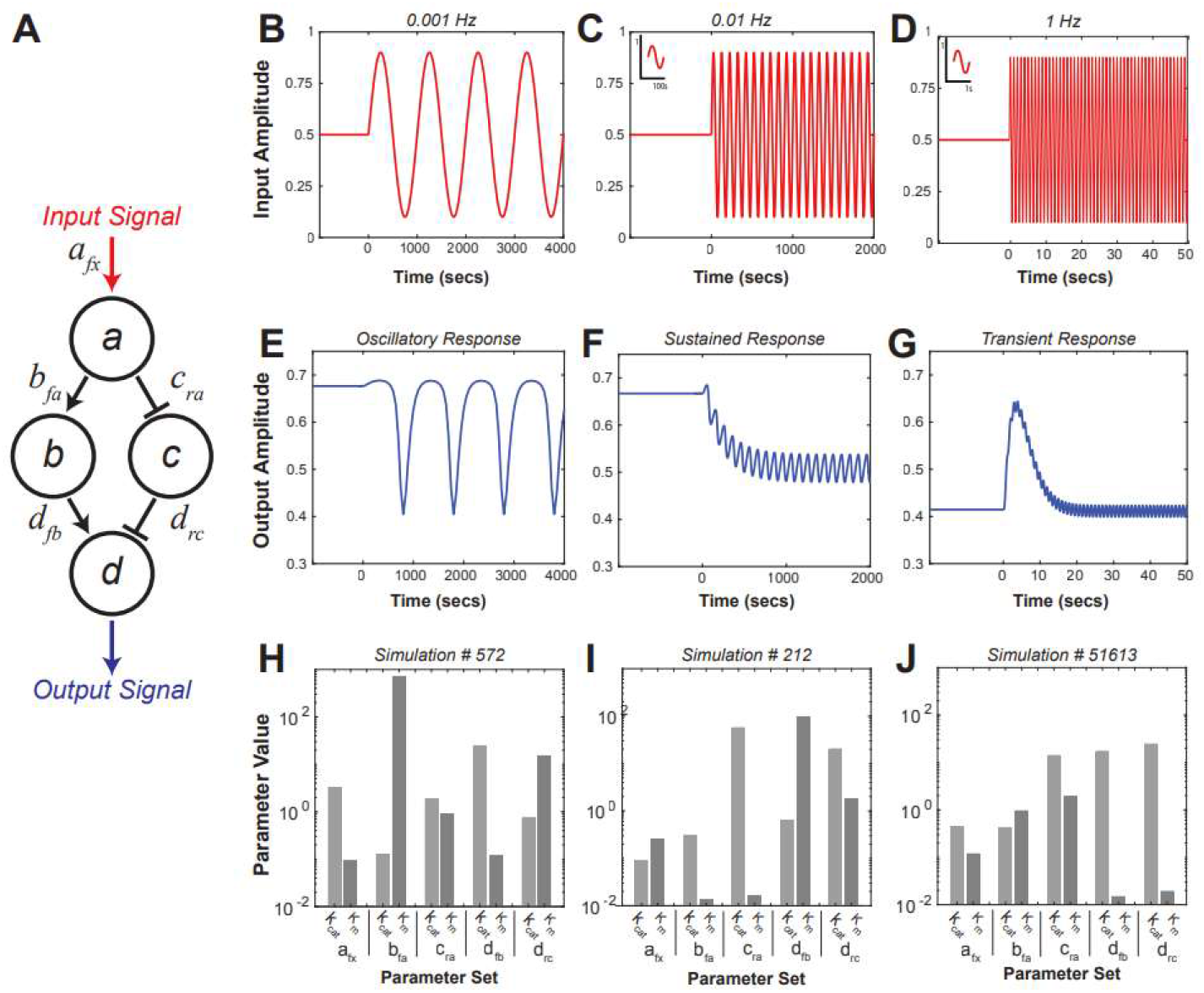
Example output responses of coherent feedforward (cFF) network, Motif 19. **(A)** Motif 19 topology. **(B-D)** Input stimuli of varying frequncies. **(E)** Oscillatory response to 0.001Hz input. **(F)** Sustained response to 0.01Hz input. **(G)** Transient response to 1Hz input. **(H-J)** Enzyme kinetic paramters for respective simulations used for further parameter analysis.

For events solved over the time vector in Equation 5, we accomplish a temporal resolution up to 2,000 times higher than the frequency-dependent time vector used for the core data. For each motif and frequency combination, 10^5^ simulations were run. Across all twenty motifs and seven frequencies, we ran 2.8×10^7^ total simulations.

After completion of all 2.8×10^7^ simulations, output data generated from each were compiled into matrices. We categorized overall motif behavior based on the characteristics of the output signal. To identify events of interests, we set forth three criteria: absolute direct current (DC) shift, decibel (dB) suppression, and maximum response. DC shift indicates the shift in the average output from the system’s steady-state prior to and after stimulus. dB suppression is an indication of how much the oscillatory amplitude was suppressed by the system after reaching steady-state following oscillatory stimulus. Maximum response gives the largest change in output amplitude following stimulus. Equations for each categorization criterion can be found in **S3 Appendix**. We chose to categorize motifs meeting such criterion into four groups based on their output characteristics, namely: oscillatory responders (**Fig 1E**), sustained responders (**Fig 1F**), and transient responders (**Fig 1G**). In order to find sustained responders, we set the criteria for absolute DC shift >0.1 and a dB suppression of > 20. In order to find transient responders, we set the criteria for absolute DC shift < 0.1, dB suppression > 20, and a maximum amplitude response > 0.2. All responses deemed to have failed the criteria for both sustained and transient response were categorized as oscillatory.

Analyzing the output responses in clusters, such as sustained and transient responders, allowed us to focus in on certain motifs and determine their sensitivities to variations in various enzyme parameters. For example, Motif 4 (*cFF*) did not produce a single sustained or transient response in any frequency. However, Motif 19 (cFF), a four-node motif, produced the most sustained responders and was one of the few motifs to produce any sort of transient response. Through this observation, additional simulations of certain motifs were run with the goal of finding additional transient responders. An additional one million simulations were run for Motifs 2 (*nFB*), 12 (*nFB/icFF*), 14 (*pFB/nFB/icFF*), 17 (*icFF*), 19 (cFF) at 1 Hz. One hertz was the chosen frequency because that was the only frequency at which a clear transient response was obtained from the initial run of 2.8×10^7^simulations.

## Results

### Motif responses can be characterized based on output characteristics

For all simulations, the output of the motif after stimulus by an oscillatory signal (**Fig 2A**) was used to characterize the motif response. An output signal can be characterized based on three characteristics, its maximum response, its sustained response, and its residual oscillations (**Fig 2B**). Most network motif responses fall under the category of noise, which do not satisfy any of our decision criteria. A select few network outputs fall under the category of sustained responders. The total number of sustained responders across all twenty motifs and all seven frequencies was 6,159; a probability occurrence percentage of 0.0022% (one sustained responder for every 5,000 simulations). Even rarer events include those categorized as transient responders. The total number of transient responders across all twenty motifs and all seven frequencies was 33, which is a probability occurrence percentage of 1.17 x 10^-4^ % (one transient responder for every million simulations).

**Fig 2.**
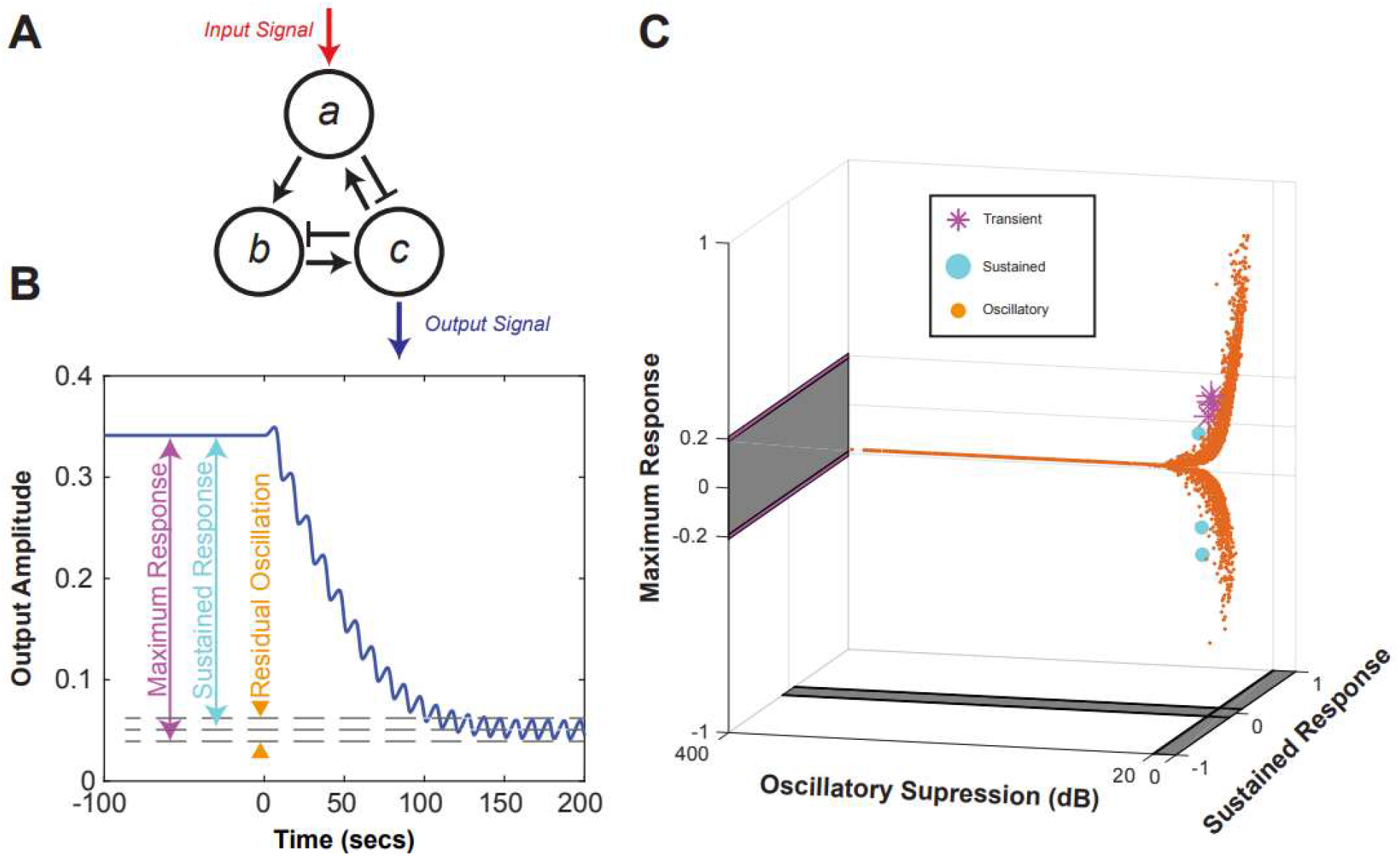
Decision criteria for output response characterization. **(A)** An example motif. The signal that is considered the output is in blue. **(B)** Definition of output characteristics through an example output signal. **(C)** All simulation responses characterized by decision criterion.

The distribution of output characteristics for each motif with responders was plotted three-dimensionally with axes of oscillatory dB suppression and maximum response showing the overall behavior of a given motif for a given frequency. Most of the simulations for all 20 motifs resulted in no change in mean output with varying grades of oscillatory suppression; furthermore, no major DC shift was observed in any of the parameter combinations that resulted in high suppression (**Fig 2C**).

### Enzymatic network motifs show median oscillatory suppression of 30-135 dBs; three-node coherent feedforward display the lowest oscillatory suppression

The average dB suppression for each motif at each frequency varied (**Fig 3**). Across all motifs, the median oscillatory suppression was the highest with an input frequency of 1Hz; median suppression increased across all seven simulated frequencies, with the exception of Motif 5 (*icFF*). Amongst all motifs, the three-node networks with coherent feedforward loops demonstrated the lowest median oscillatory suppression, suggesting that such feedforward loops act as poorer suppressors of noise relative to other network topologies. In contrast, Motif 16 (*icFF*), demonstrated the highest propensity to suppress the oscillatory input signal. Other motifs with similarly high propensity to suppress oscillatory signals were Motif 6 (*pFBx2*), Motif 7 (*nFBx2*), and Motif 8 (*pFB/nFB*).

**Fig 3.**
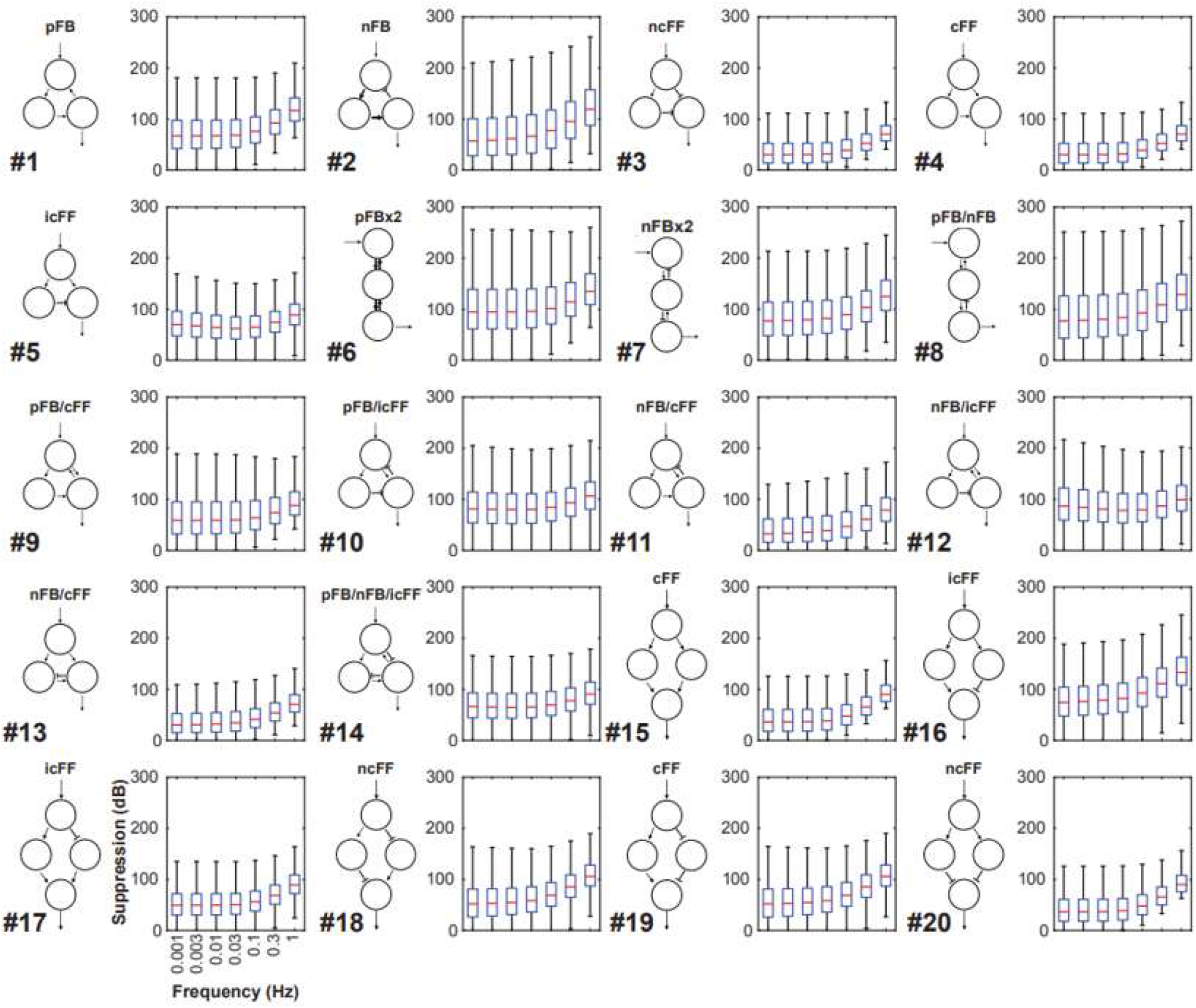
Median oscillatory suppression across all simulation for each motif.

The occurrence probabilities for oscillatory suppression with varying magnitude of attenuation was calculated across all 20 motifs and simulated frequencies (**S4 Fig**). In line with trends of median dB suppression, higher frequency inputs were more likely to be suppressed across all motifs. Some motifs, particularly those with incoherent feedforward (*icFF*), produce similar suppression across all frequencies. Other motifs, particularly those with coherent feedforward (cFF) and negative-coherent feedforward (*ncFF*), produced much higher suppression in higher frequencies than in lower frequencies.

### Input-output responses depend on network topology and stimulus frequency

The distribution of network topologies that transduced sustained (**Fig 4A**) and transient **(Fig 4B**) responses was not evenly dispersed across the 20 network motifs. Most motifs induced minimal change onto the input signal, whereas others transduced sustained and/or transient responses when stimulated by signals at certain characteristic frequencies. Some motifs outputted sustained and/or transient responses at much higher rates than others. Motifs that were particularly likely to transduce sustained responses all had nodes with coherent feedforward (Motifs 18, 19, 11) and/or negative feedback (Motifs 2, 11). In addition, motifs most likely to produce transient responses both contained nodes with incoherent feedforward (Motifs 14, 17). No motifs were capable of transducing sustained or transient responses given the entire range of stimulus frequencies. Additionally, several motifs could shift the DC output of the system in response to oscillatory input. The probability occurrence of varying magnitudes of absolute DC shift transduced by each motif at each frequency was found to vary across network topologies and frequencies, with lower frequencies being the most likely to transduce higher absolute DC shift across all motifs (**S5 Fig**).

**Fig 4.**
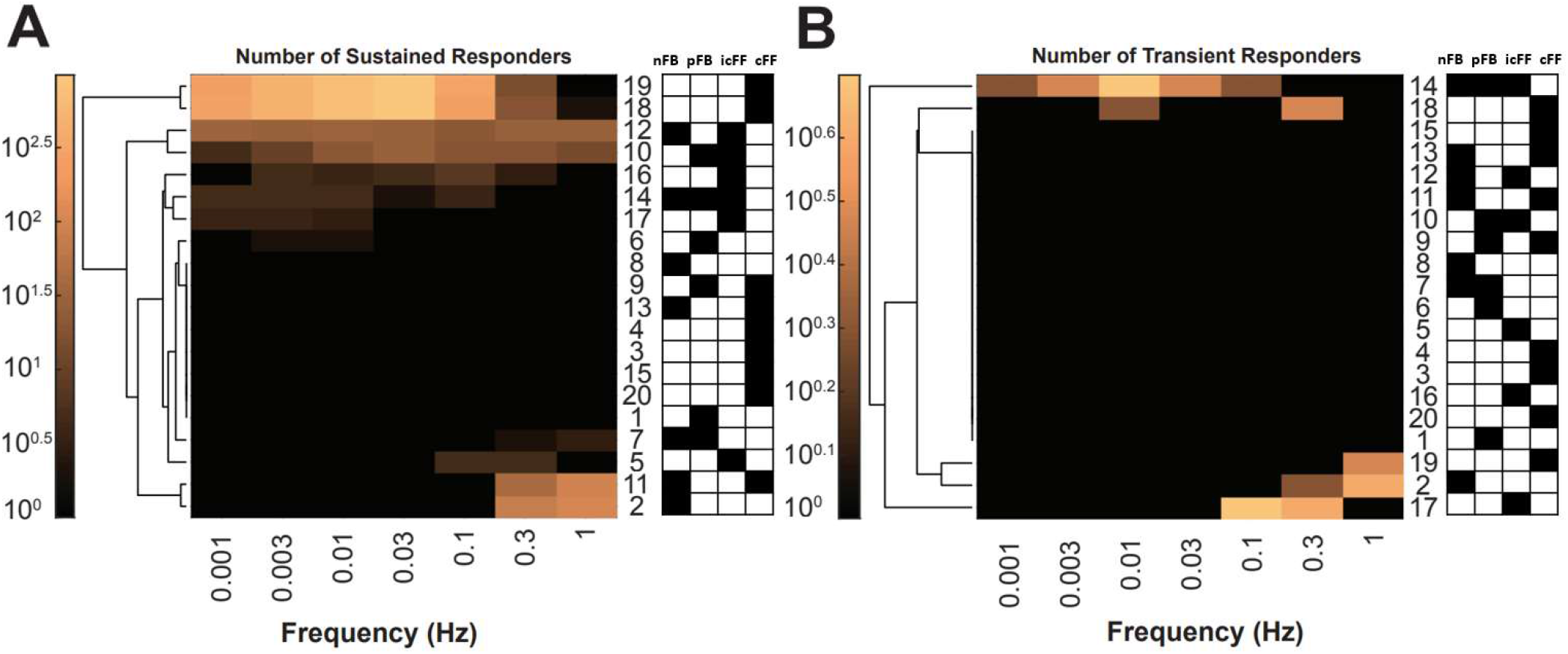
Number of responders for each motif at all frequencies. (**A**) Number of sustained responders. (**B**) Number of transient responders.

### Identifying potential mechanisms for oscillatory signal processing using parameter sensitivity analysis of incoherent feedforward motifs

It was evident that certain motifs are capable of behaving as sustained and transient responders at specific frequencies. In particular, Motif 19 (*cFF*) was identified as a network topology with a high percentage of both sustained and transient responders relative to other topologies.

In order to investigate the metrics of the network that may be responsible for such output behavior, we followed up our initial simulations with a more highly resolved parameter sweep for Motif 19 (cFF). We ran a total of 10^5^ additional simulations across all seven frequencies for Motif 19. Out of the 10^5^ additional simulations of Motif 19, there were a total of 2,201 transient responders (8 at 0.1 Hz; 34 at 0.3 Hz; 2,159 at 1 Hz), a transient response probability occurrence of 2.2%. Transient responders at 1 Hz greatly outnumber those at lower frequencies, even within this same network topology.

We also ran an additional transient sweep across all parameters, to see if we could transduce additional sustained and transient responses. The parameter sets that induced sustained responses were recorded and compared to an equal number of randomly selected parameters that did not induce sustained responses. It was found that Motif 19 (cFF) can output a DC shifted, sustained output at all simulated frequencies. A representative selection of negatively shifted sustained outputs of Motif 19 at input frequencies of 0.001, 0.003, 0.01, 0.03, and 0.1, and 0.3Hz are shown in **Fig 5A**. We note that different time scales are shown to retain detail at varying frequencies. Comparison of parameters for these sustained responders (**Fig 5B**) versus randomly selected equal number of non-responders (**Fig 5C**) show that only a small number of parameter values are adequate to achieve sustained response across input frequencies. Due to the Law of Large Numbers, the median value for each parameter in **Fig 5C** approaches the mean of our logarithmic parameter sweep. In contrast to the randomly selected parameters, we see that the median parameter values of networks that transduce sustained responses stray from the median value of the logarithmic sweep. This indicates that enzyme parameters, within the same network topology, impact network response to oscillatory information.

**Fig 5.**
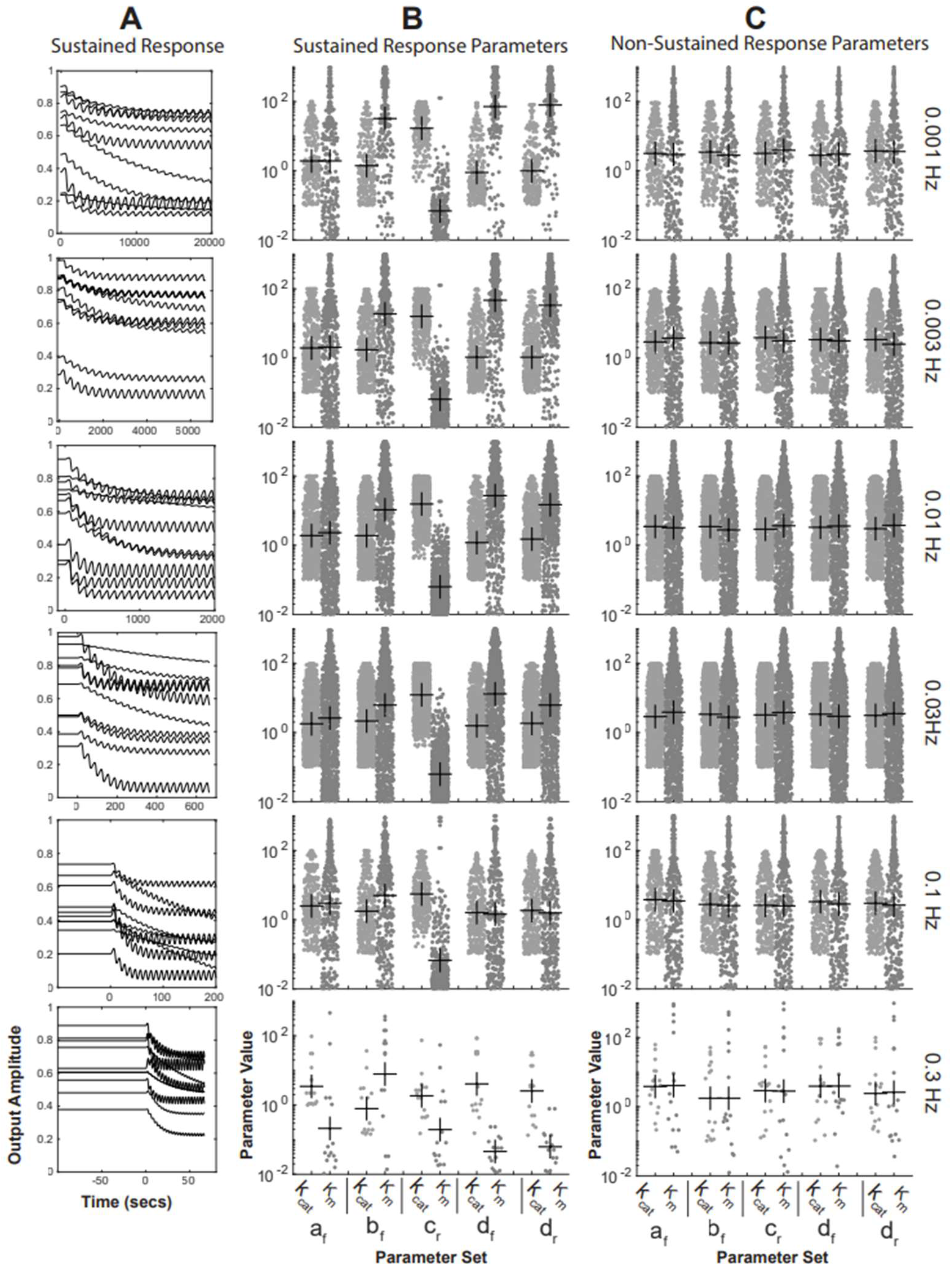
Comparison of enzyme parameters of Motif 19 which produced negative-shifted sustained responses. **(A)** Negative-shifted sustained output responses transduced through Motif 19 at various frequencies. **(B)** Enzyme parameters of Motif 19 which produce each sustained response in (A). **(C)** An equal number of non-sustained response producing parameter sets selected from Motif 19 simulations.

Of note, the topology governed the sensitivity of different enzymatic parameters (**Fig 5B**). In **Fig 5B**, the density of the parameter values, which produce sustained outputs, around the median reflects how resilient the topology is to variation of that parameter. A larger spread of values indicates higher resiliency to parameter variation, whereas a high density of parameters around the median suggests that the parameter value must be very close to the median to still transduce sustained responses. Unsurprisingly, at higher frequencies, enzyme parameters across all nodes demonstrated less resilience to change. Interestingly, *K_m_* of node *c_ra_*, the node inhibited by *a* and that inhibits d, was the least resilient parameter across all frequencies. The *K_m_* of this node needed to be quite small (<10^-1^ mM) to produce a sustained output. In contrast, the *K_m_* of node *b_ra_* had to be relatively high (>10mM) to produce a transient sustained. Identical analysis of sustained responders with positive shifts demonstrated similar trends across node parameters (**S6 Fig**).

In addition to the sustained response parameter sweep on Motif 19 (*cFF*), we ran additional parameter analysis to determine sensitivity to parameters in the transduction of transient responses. In this analysis, for every 10^4^ simulations, nine parameters were held constant, and a random parameter value was assigned to the one nonfixed parameter. The constant values used were the same parameter values for both the transient output in **Fig 1G** and **1J**. These constant values were chosen because we found that they produced very sharp transient responses in our initial simulation batch **(Fig 1G).** Parameter values that maintained similar transient behavior when all others were held constant were recorded to gauge system sensitivity for each network parameter. Parameter sweeping across the transient outputs of Motif 19 (*cFF*) also indicated varying levels of sensitivity to different network parameters (**S7 Fig**). This analysis demonstrated that transient response was very sensitive to the enzymatic parameter combinations of nodes A, B, and C. *K_m,drc_* and *K_m,dfb_* were found to be the most resilient parameters for transduction of transient behavior given an oscillatory input stimulus. Thus, node D’s *K_m_* value was deemed less responsible for the transient behavior compared to the other nodes. It is also interesting to compare the relative values of parameters that transduce transient behavior to those that transduce sustained behavior within Motif 19. For example, in the transduction of sustained responses, Michaelis constant *K_m,cra_* was very low. In contrast, very high *K_m,cra_* values were necessary in the transduction of a transient response.

### Motif 12 is capable of behaving as a perpetual oscillator and as a frequency modulator

In **Fig 6**, we see response characteristics unique to Motif 12 (*nFB/icFF*). Amongst all simulations, only Motif 12 exhibited oscillatory behavior at equilibrium (**Fig 6A, B**). Additionally, Motif 12 is the only network topology for which, given specific parameters, the system response is resistant to the input signal (**Fig 6A**). This is evident by the fact that Motif 12 can produce an oscillatory output at equilibrium and continues to produce an oscillatory output after application of a stimulus, thereby behaving mostly independent of the input signal. The network, in such cases, maintains the central frequency of its equilibrium oscillation despite the introduction of an input signal at a different frequency. However, in these cases, the output of the system exhibits beat-like behavior, with the amplitude of the output varying significantly at a lower frequency similar to that of the input stimulus.

**Fig 6.**
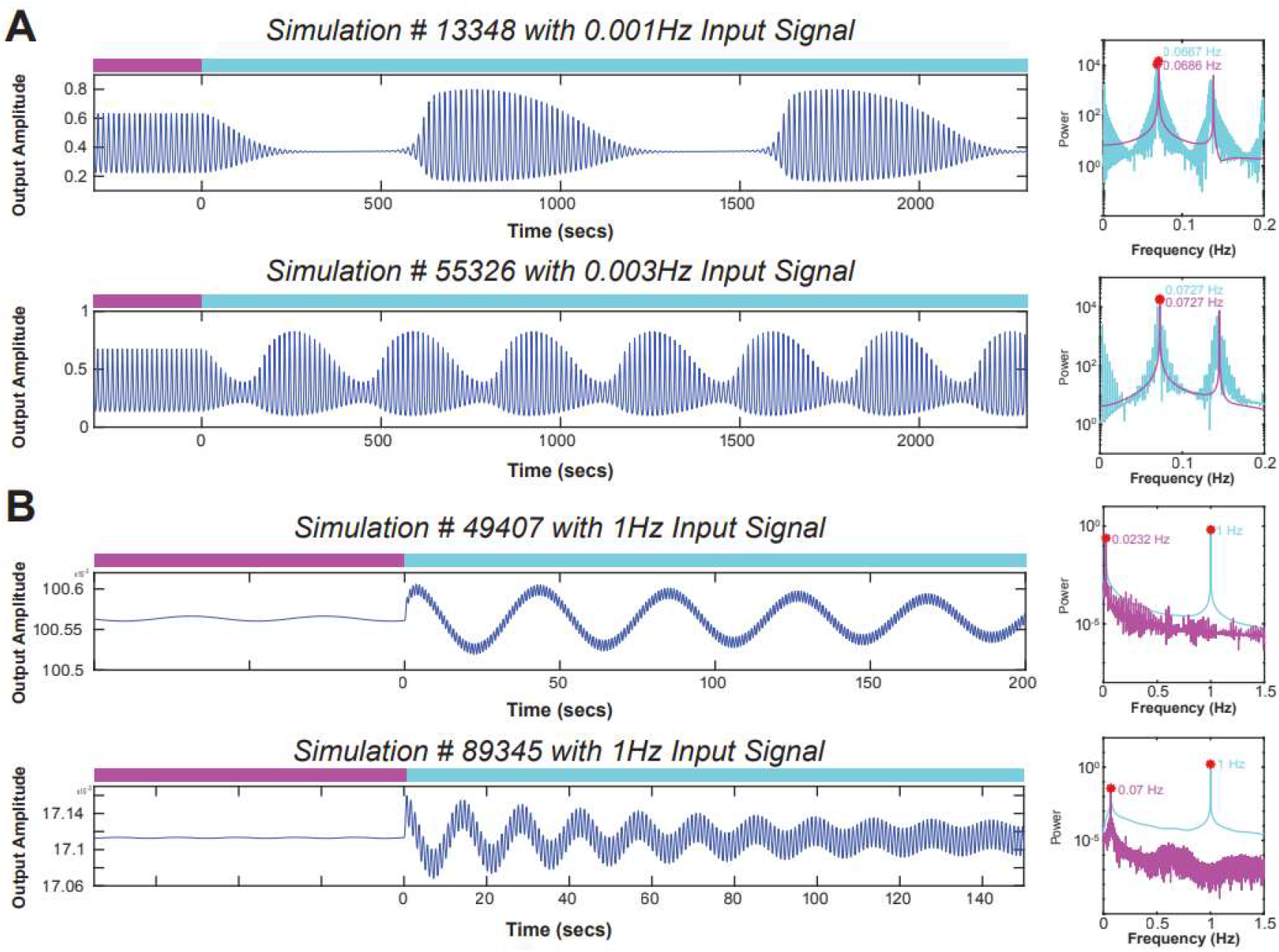
Perpetual oscillatory responses of Motif 12. **(A)** Oscillatory responses of Motif 12 which are independent of input stimulus. **(B)** Frequency modulation of equilibrium oscillation by transduction of an oscillatory input signal through Motif 12.

Alternatively, certain configurations of Motif 12 (*nFB/icFF*) exhibit the ability to behave as frequency modulators given sinusoidal stimulus. We have discussed that Motif 12, under equilibrium, oscillates at varying frequencies depending on enzyme parameters. Two such examples of frequency modulation due to stimulus are shown in **Figure 6B**. The first example behaves as a steady-state oscillator in equilibrium, oscillating at a frequency of 0.0232Hz. Interestingly, after stimulus by a 1Hz input, the system outputs a waveform with a central frequency of exactly 1Hz, which is a 43X decrease in the cyclic period of the system output. Similarly, a network with different parameters, which oscillated in equilibrium at 0.07Hz, also responded to a 1Hz input with a frequency modulation, outputting a sinusoid at a 1Hz frequency after stimulus, a 14.3X decrease in the output’s period. Such drastic frequency modulations of network outputs have the potential to greatly impact a biological network.

Furthermore, a Motif 12 network with the same parameter set, which acts as a perpetual oscillator at equilibrium, demonstrates different behavior at different stimulus frequencies. After input stimulus of 0.001Hz, the system acts as a simple sustained responder. In contrast, after a stimulus of 0.1Hz, the same motif becomes a frequency modulator.

## Discussion

Complex biological systems in nature and their subcomponents often evolve to enrich for certain behaviors that are advantageous for survival or reproduction. Considering the cyclic nature of most physical phenomena, it is logical to presume that cell biological signaling networks are tuned to handle information that operates at certain characteristic frequencies. In our computational paradigm, we prescribed a net zero input (ranging from normalized −0.5 to 0.5 with a characteristic frequency) into 20 commonly observed network motifs, which were perturbed in their steady state equilibria. The expansive non-zero input-output relationships explored in this study demonstrates that only certain motifs and motif combinations can reliably process oscillatory signals while most others act as noise suppressors. This behavior has likely evolved to preserve the robustness of cell biological operations that are critical for physiological function.^20^

Our results show that enzymatic network motifs can decode information from oscillatory input signals in a frequency- and enzyme parameter-dependent manner. Importantly, we observed that enzymatic networks that are natural resonators can use the inherent robustness of negative feedback loops to suppress noise from input signals with characteristic frequencies that differ from the tuned frequency of the network.

We further show that networks can produce responses to oscillatory input beyond noise suppression, outputs which were categorized either as transient or sustained responses. Response type and severity were dependent on both the parameters of the network and the frequency of the input stimulus. Additionally, negative feedback and four-node feedforward topologies were shown to act as contextual switches under oscillatory inputs, both as frequency modulators and AC/DC converters. The frequency modulating behavior of Motif 12 (*nFB/icFF*), in particular, is interesting as it relates to allosteric inhibition within biochemical networks. Frequency modulation is an important product of inhibitory networks, as such networks regulate many cellular processes such as the rates of glycolysis, growth, regeneration, and more.^23,24,25,26^ It may be worth to further follow up which network parameters are most vital to frequency modulation and to gain better understanding of the resilience of such regulation networks as they relate to diseases, such as cancer.

Some of the motifs, such as bifans, could produce consistent steady state outputs even when the input is an oscillatory fluctuation with a net mean change of zero from the initial condition. Bifans motifs that are overrepresented especially in transcriptional networks and are known to act as noise filters and coincidence detectors under certain logic conditions.^21^ As such, it is unsurprising that they could act as transient signal transducers in the face of oscillatory input signals.

Certain motifs were shown to transduce both transient and sustained responses depending on their system parameters. Most notably, Motif 19, a four-node network with coherent feedforward, was identified as a likely network motif to produce either transient or sustained outputs given an oscillatory input stimulus. Parameter analysis of Motif 19 (*cFF*) showed that output type is sensitive to the enzymatic parameters; however, the output responses of Motif 19 under all frequencies were most impacted by the changes in the inhibitory reaction disassociation constant, *K_m,ca_*, and the intermediate forward reaction disassociation constant, *K_m,ba_* in the transduction of sustained responses. For transduction of transient responses in Motif 19, all parameters except *K_m,drc_* and *K_m,drb_* were highly sensitive. For several nodes, the relative enzymatic values that produced a sustained response were different than those which produced a transient response. These results indicate that the relative values of enzyme parameters compared to others within a network are likely to play a role in regulating oscillatory signal transduction. Further follow up is necessary to evaluate whether relative ratios among parameters or the absolute value of a given parameter is more critical for overall network behavior.

In conclusion, we show that biologically relevant network motifs are capable of processing and decoding information from oscillatory input signals. We show the importance of network topology, enzyme kinetics, and signal frequency on the processing of oscillatory cellular information. In addition to demonstrating varying sensitivity to different enzyme parameters, we show that network output depended on the frequency of the input signal. Even within Motif 19, which was most likely to produce sustained outputs, enzyme parameters had to be very finely tuned to produce any sustained output at certain frequencies. However, at other frequencies, the system was more resilient to variations in enzyme kinetics. These results demonstrate the importance of input signal frequency to the transduction of information through an enzymatic network. Enzyme networks appear to be tuned to transduce signals with certain characteristic frequencies while suppressing signals with frequencies that do not match the tuning of the system. Network tuning to specific signal frequencies could be a mechanism in which different pathways transduce proper signals while ignoring noise from other cellular pathways.

## Supporting information

S2 Appendix - Motif ODEs

S3 Appendix - Output Characteristic Equations

All Supplementary Figures

## Acknowledgements

The project was supported by the NIH (R01DK118222 and R01DK131047). TKF was in part supported by R25DK124917.

## Data and Code Availability

All the data supporting the findings of this study are available within the manuscript and its supplementary information. The annotated Matlab code, all of the simulation parameters and the raw data presented here can be found in our GitHub page https://github.com/AzelogluLab/Oscillatory-Motifs.

## Authorship Contributions

J. Rejas: Conceptualization, Programming, Performed Simulations, Investigation, Analysis Methodology, Writing – Initial Draft

T. Fallon: Analysis, Visualization, Performed Simulations, Writing – Initial Draft, Writing – Editing

A. M. Leader: Programming, Performed Simulations, Analysis, Visualization, Writing – Initial Draft

E. U. Azeloglu: Conceptualization, Funding acquisition, Investigation, Project administration, Resources, Supervision,Visualization, Writing – review & editing

