## Supplementary material for "Biochemical network motifs can transduce and process oscillatory information": S2 Appendix - Motif ODEs

**S2 Appendix.** Systems of ODEs used to represent each network motif.

Motif #1: Positive Feedback (pFB)

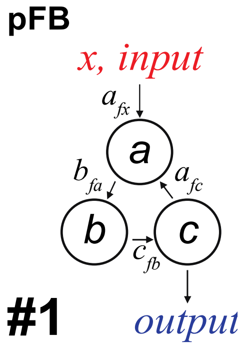


$$\frac{da}{dt} = \frac{(k_{cat_{afx}}\boldsymbol{x}(1-a))}{(K_{m_{afx}} + (1-a))} + \frac{(k_{cat_{afc}}(c*(1-a)))}{(K_{m_{afc}} + (1-a))} - \frac{(k_{cat_{ar}} * 0.5 * a)}{(K_{m_{ar}} + a)}$$

$$\frac{db}{dt} = \frac{(k_{cat_{bfa}}(a*(1-b)))}{(K_{m_{bf}} + (1-b))} - \frac{(k_{cat_{br}} * 0.5 * b)}{(K_{m_{br}} + b)}$$

$$\frac{dc}{dt} = \frac{(k_{cat_{cfb}}(b*(1-c)))}{(K_{m_{cfb}} + 1 - c)} - \frac{(k_{cat_{cr}} * 0.5 * c)}{(K_{m_{cr}} + c)}$$

---

Motif #2: Negative Feedback (nFB)

$$\frac{da}{dt} = \frac{(k_{cat_{afx}}\boldsymbol{x}(1-a))}{(K_{m_{afx}} + (1-a))} - \frac{(k_{cat_{arc}} * c * a)}{(K_{m_{arc}} + a)}$$

$$\frac{db}{dt} = \frac{(k_{cat_{bfa}}(a*(1-b)))}{(K_{m_{bfa}} + (1-b))} - \frac{(k_{cat_{br}} * 0.5 * b)}{(K_{m_{br}} + b)}$$

$$\frac{dc}{dt} = \frac{(k_{cat_{cfb}}(b*(1-c)))}{(K_{m_{cfb}} + (1-c))} - \frac{(k_{cat_{cr}} * 0.5 * c)}{(K_{m_{cr}} + c)}$$

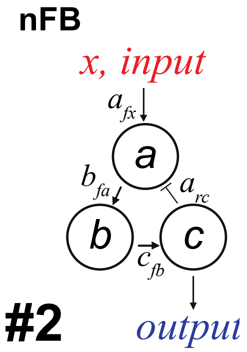


---

Motif # 3: Negative Coherent Feedforward (ncFF)

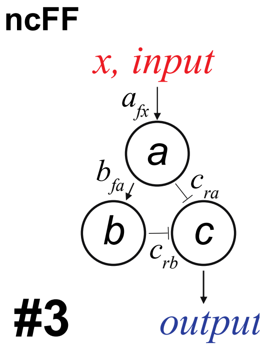


$$\frac{da}{dt} = \frac{(k_{cat_{afx}}\boldsymbol{x}(1-a))}{(K_{m_{afx}} + (1-a))} - \frac{(k_{cat_{ar}} * 0.5 * a)}{(K_{m_{ar}} + a)}$$

$$\frac{db}{dt} = \frac{(k_{cat_{bfa}}(a*(1-b)))}{(K_{m_{bfa}} + (1-b))} - \frac{(k_{cat_{br}} * 0.5 * b)}{(K_{m_{br}} + b)}$$

$$\frac{dc}{dt} = \frac{(k_{cat_{cf}} * 0.5 * (1-c))}{(K_{m_{cf}} + 1 - c)} - \frac{(k_{cat_{cra}} * a * c)}{(K_{m_{cra}} + c)} - \frac{(k_{cat_{crb}} * b * c)}{(K_{m_{crb}} + c)}$$

---

Motif #4: Coherent Feedforward (cFF)

$$\frac{da}{dt} = \frac{(k_{cat_{afx}}\boldsymbol{x}(1-a)}{(K_{m_{afx}} + (1-a))} - \frac{(k_{cat_{ar}} * 0.5 * a)}{(K_{m_{ar}} + a)}$$

$$\frac{db}{dt} = \frac{(k_{cat_{bfa}}(a * (1-b))}{(K_{m_{bfa}} + (1-b))} - \frac{(k_{cat_{br}} * 0.5 * b)}{(K_{m_{br}} + b)}$$

$$\frac{dc}{dt} = \frac{(k_{cat_{cfa}} * a * (1-c)))}{(K_{m_{cfa}} + (1-c)))} + \frac{(k_{cat_{cfb}} * b * (1-b)))}{(K_{m_{cfb}} + (1-c)))} - \frac{(k_{cat_{cr}} * 0.5 * c))}{(K_{m_{cr}} + c))}$$

**cFF**

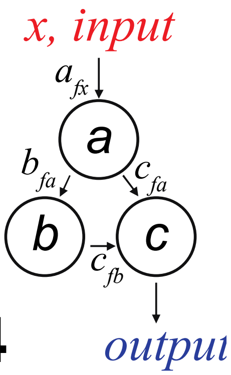


**#4**

---

Motif #5: Incoherent Feedforward (icFF)

**icFF**

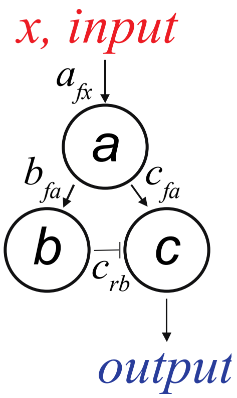


**#5**

$$\frac{da}{dt} = \frac{(k_{cat_{afx}}\boldsymbol{x}(1-a)}{(K_{m_{afx}} + (1-a))} - \frac{(k_{cat_{ar}} * 0.5 * a)}{(K_{m_{ar}} + a)}$$

$$\frac{db}{dt} = \frac{(k_{cat_{bfa}}(a * (1-b))}{(K_{m_{bfa}} + (1-b))} - \frac{(k_{cat_{br}} * 0.5 * b)}{(K_{m_{br}} + b)}$$

$$\frac{dc}{dt} = \frac{(k_{cat_{cfa}} * b * (1-c))}{(K_{m_{cfa}} + 1 - c)} - \frac{(k_{cat_{crb}} * b * a)}{(K_{m_{crb}} + c)}$$

---

Motif #6: Positive Feedback x2 (pFB x2)

$$\frac{da}{dt} = \frac{(k_{cat_{afx}}\boldsymbol{x}(1-a)}{(K_{m_{afx}}+1-a)} + \frac{(k_{cat_{afb}}(b*(1-a))}{(K_{m_{afb}}+1-a)} - \frac{(k_{cat_{ar}}*0.5*a)}{(K_{m_{ar}}+a)}$$

**pFBx2**

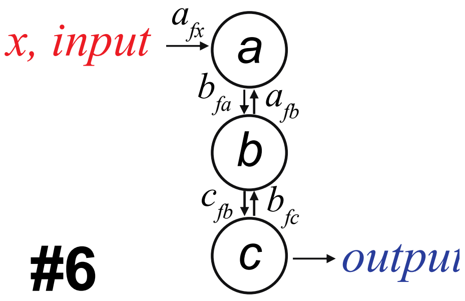


**#6**

$$\frac{db}{dt} = \frac{(k_{cat_{bfa}}*a*(1-b)}{(K_{m_{bf\boldsymbol{a}}}+1-b)} + \frac{(k_{cat_{bfc}}(c*(1-b))}{(K_{m_{bfc}}+1-b)} - \frac{(k_{cat_{br}}*0.5*b)}{(K_{m_{br}}+b)}$$

$$\frac{dc}{dt} = \frac{(k_{cat_{cfb}}*b*(1-c)}{(K_{m_{cfb}}+1-c)} - \frac{(k_{cat_{cr}}*0.5*c)}{(K_{m_{cr}}+c)}$$

---

Motif #7: Positive Feedback/Negative Feedback (pFB/nFB)

**pFB/nFB**

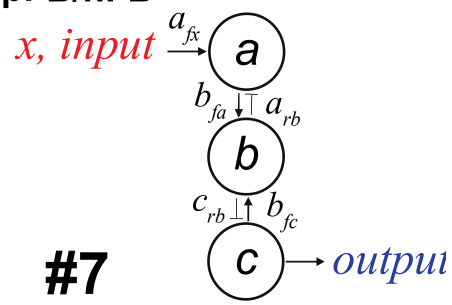


**#7**

$$\frac{da}{dt} = \frac{(k_{cat_{afx}}\boldsymbol{x}(1-a)}{(K_{m_{afx}}+1-a)} - \frac{(k_{cat_{arb}}(b*a))}{(K_{m_{arb}}+a)}$$

$$\frac{db}{dt} = \frac{(k_{cat_{bfa}}(a*(1-b))}{(K_{m_{bfa}}+1-b))} + \frac{(k_{cat_{bfc}}*c*1-b))}{(K_{m_{bfc}}+1-b))} - \frac{(k_{cat_{br}}*0.5*b))}{(K_{m_{br}}+b))}$$

$$\frac{dc}{dt} = \frac{(k_{cat_{cf}}(0.5*(1-c))}{(K_{m_{cf}}+(1-c))} - \frac{(k_{cat_{crb}}*b*c)}{(K_{m_{crb}}+c)}$$

---

Motif #8: Positive Feedback/Negative Feedback???

$$\frac{da}{dt} = \frac{\left(k_{cat_{afx}} \boldsymbol{x}(1-a)\right)}{\left(K_{m_{afx}} + 1 - a\right)} + \frac{\left(k_{cat_{afb}}\left(b * (1-a)\right)\right)}{\left(K_{m_{afb}} + 1 - a\right)} - \frac{\left(k_{cat_{ar}} * 0.5 * a\right)}{\left(K_{m_{ar}} + a\right)}$$

$$\frac{db}{dt} = \frac{\left(k_{cat_{bfa}}\left(a * (1-b)\right)\right)}{\left(K_{m_{bfa}} + 1 - b\right)} - \frac{\left(k_{cat_{brc}} * c * \mathrm{b}\right)}{\left(K_{m_{brc}} + b\right)}$$

$$\frac{dc}{dt} = \frac{\left(k_{cat_{cfb}} * b * (1-c)\right)}{\left(K_{m_{cfb}} + 1 - c\right)} - \frac{\left(k_{cat_{cr}} * 0.5 * c\right)}{\left(K_{m_{cr}} + c\right)}$$

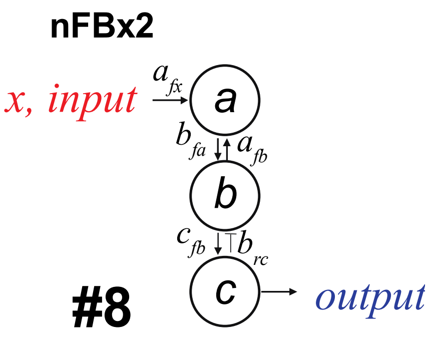


---

Motif #9: Positive Feedback/Coherent Feedforward (pFB/cFF)

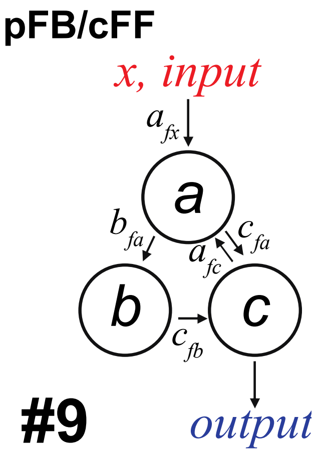


$$\frac{da}{dt} = \frac{\left(k_{cat_{afx}} \boldsymbol{x}(1-a)\right)}{\left(K_{m_{afx}} + 1 - a\right)} + \frac{\left(k_{cat_{afc}}\left(c * (1-a)\right)\right)}{\left(K_{m_{afc}} + 1 - a\right)} - \frac{\left(k_{cat_{ar}} * 0.5 * a\right)}{\left(K_{m_{ar}} + a\right)}$$

$$\frac{db}{dt} = \frac{\left(k_{cat_{bfa}}\left(a * (1-b)\right)\right)}{\left(K_{m_{bfa}} + 1 - b\right)} - \frac{\left(k_{cat_{br}} * 0.5 * b\right)}{\left(K_{m_{br}} + b\right)}$$

$$\frac{dc}{dt} = \frac{\left(k_{cat_{cfa}} * a * (1-c)\right)}{\left(K_{m_{cfa}} + 1 - c\right)} + \frac{\left(k_{cat_{cfb}} * b * (1-c)\right)}{\left(K_{m_{cfb}} + 1 - c\right)} - \frac{\left(k_{cat_{cr}} * 0.5 * c\right)}{\left(K_{m_{cr}} + c\right)}$$

---

Motif #10: Positive Feedback/Incoherent Feedforward (pFB/icFF)

$$\frac{da}{dt} = \frac{(k_{cat_{afx}} \boldsymbol{x}(1-a)}{(K_{m_{afx}} + 1 - a))} - \frac{(k_{cat_{arc}} * c * a))}{(K_{m_{arc}} + a))}$$

$$\frac{db}{dt} = \frac{(k_{cat_{bfa}}(a * (1-b))}{(K_{m_{bfa}} + 1 - b))} - \frac{(k_{cat_{br}} * 0.5 * b))}{(K_{m_{br}} + b))}$$

$$\frac{dc}{dt} = \frac{(k_{cat_{cfa}} * a * (1-c))}{(K_{m_{cfa}} + 1 - c))} - \frac{(k_{cat_{crb}} * b * c)}{(K_{m_{crb}} + c)}$$

**pFB/icFF**

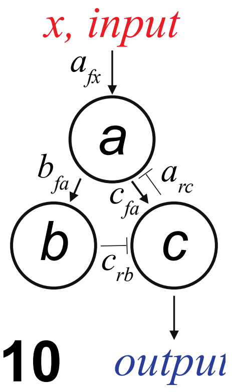


**#10**

---

Motif #11: Negative Feedback/Coherent Feedforward (nFB/cFF)

**nFB/cFF**

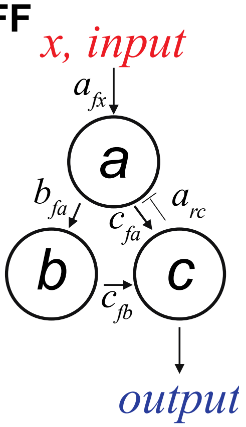


**#11**

$$\frac{da}{dt} = \frac{(k_{cat_{afx}} \boldsymbol{x}(1-a)}{(K_{m_{afx}} + 1 - a))} - \frac{(k_{cat_{arc}} * c * a)}{(K_{m_{arc}} + a)}$$

$$\frac{db}{dt} = \frac{(k_{cat_{bfa}}(a * (1-b))}{(K_{m_{bfa}} + 1 - b)} - \frac{(k_{cat_{br}} * 0.5 * b)}{(K_{m_{br}} + b)}$$

$$\frac{dc}{dt} = \frac{(k_{cat_{cfa}} * a * (1-c))}{(K_{m_{cfa}} + 1 - c))} + \frac{(k_{cat_{cfb}} * b * (1-c))}{(K_{m_{cfb}} + 1 - c))} - \frac{(k_{cat_{cr}} * 0.5 * c))}{(K_{m_{cr}} + c))}$$

---

Motif #12: Negative Feedback/Incoherent Feedforward (nFB/icFF)

$$\frac{da}{dt} = \frac{(k_{cat_{afx}}\boldsymbol{x}(1-a)}{(K_{m_{afx}}+1-a)} + \frac{(k_{cat_{afc}}(c*(1-a))}{(K_{m_{afc}}+1-a)} - \frac{(k_{cat_{ar}}*0.5*a)}{(K_{m_{ar}}+a)}$$

$$\frac{db}{dt} = \frac{(k_{cat_{bfa}}(a*(1-b))}{(K_{m_{bfa}}+1-b)} - \frac{(k_{cat_{br}}*0.5*b)}{(K_{m_{br}}+b)}$$

$$\frac{dc}{dt} = \frac{(k_{cat_{cfa}}(a*(1-c))}{(K_{m_{cfa}}+1-c)} - \frac{(k_{cat_{crb}}*0.5*b*c)}{(K_{m_{crb}}+c)}$$

**nFB/icFF**

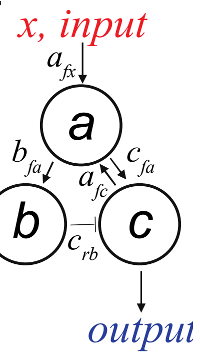


**#12**

---

Motif #13: Negative Feedback/Coherent Feedforward (nFB/cFF)

**nFB/cFF**

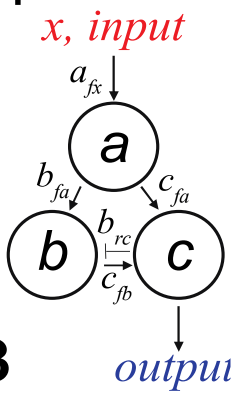


**#13**

$$\frac{da}{dt} = \frac{(k_{cat_{afx}}\boldsymbol{x}(1-a)}{(K_{m_{afx}}+1-a)} - \frac{(k_{cat_{ar}}*0.5*a)}{(K_{m_{ar}}+a)}$$

$$\frac{db}{dt} = \frac{(k_{cat_{bfa}}(a*(1-b))}{(K_{m_{bfa}}+1-b))} - \frac{(k_{cat_{brc}}*c*\mathrm{b})}{(K_{m_{brc}}+b)}$$

$$\frac{dc}{dt} = \frac{(k_{cat_{cfa}}*a*(1-c))}{(K_{m_{cfa}}+1-c))} + \frac{(k_{cat_{cfb}}*b*(1-c))}{(K_{m_{cfb}}+1-c))} - \frac{(k_{cat_{cr}}*0.5*c))}{(K_{m_{cr}}+c))}$$

---

Motif #14: Positive Feedback/Negative Feedback/Incoherent Feedforward (pFB/nFB/icFF)

$$\frac{da}{dt} = \frac{(k_{cat_{afx}}\boldsymbol{x}(1-a)}{(K_{m_{afx}}+1-a)} + \frac{(k_{cat_{afc}}(c*(1-a))}{(K_{m_{afc}}+1-a))} - \frac{(k_{cat_{ar}}*0.5*a))}{(K_{m_{ar}}+a))}$$

$$\frac{db}{dt} = \frac{(k_{cat_{bfa}}(a*(1-b))}{(K_{m_{bfa}}+1-b))} - \frac{(k_{cat_{brc}}*c*b))}{(K_{m_{brc}}+b))}$$

$$\frac{dc}{dt} = \frac{(k_{cat_{cfb}}(b*(1-c))}{(K_{m_{cfb}}+1-c))} - \frac{(k_{cat_{cra}}*a*c))}{(K_{m_{cra}}+c)}$$

**pFB/nFB/icFF**

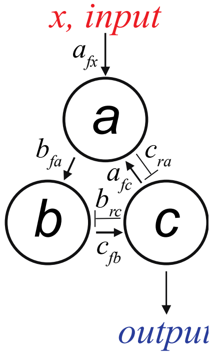


**#14**

---

Motif #15: Coherent Feedforward (cFF)

**cFF**

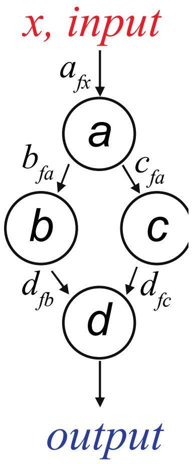


**#15**

$$\frac{da}{dt} = \frac{(k_{cat_{afx}}\boldsymbol{x}(1-a)}{(K_{m_{afx}}+1-a))} - \frac{(k_{cat_{ar}}(0.5*a))}{(K_{m_{ar}}+a))}$$

$$\frac{db}{dt} = \frac{(k_{cat_{bfa}}(a*(1-b))}{(K_{m_{bfa}}+1-b))} - \frac{(k_{cat_{br}}*0.5*b))}{(K_{m_{br}}+b))}$$

$$\frac{dc}{dt} = \frac{(k_{cat_{cfa}}(a*(1-c))}{(K_{m_{cfa}}+1-c))} - \frac{(k_{cat_{cr}}*0.5*c))}{(K_{m_{cr}}+c)}$$

$$\frac{dd}{dt} = \frac{(k_{cat_{dfb}}(b*(1-d))}{(K_{m_{dfb}}+1-d))} + \frac{(k_{cat_{dfc}}(c)*(1-d))}{(K_{m_{dfc}}+1-d))} - \frac{(k_{cat_{dr}}*0.5*d))}{(K_{m_{dr}}+d))}$$

---

Motif #16: Incoherent Feedforward (icFF)

$$\frac{da}{dt} = \frac{(k_{cat_{afx}}\boldsymbol{x}(1-a)}{(K_{m_{afx}}+1-a))} - \frac{(k_{cat_{ar}}(0.5*a))}{(K_{m_{ar}}+a))}$$

$$\frac{db}{dt} = \frac{(k_{cat_{bfa}}(a*(1-b))}{(K_{m_{bfa}}+1-b))} - \frac{(k_{cat_{br}}*0.5*b))}{(K_{m_{br}}+b))}$$

$$\frac{dc}{dt} = \frac{(k_{cat_{cfa}}(a*(1-c))}{(K_{m_{cfa}}+1-c)} - \frac{(k_{cat_{cr}}*0.5*c)}{(K_{m_{cr}}+c)}$$

$$\frac{dd}{dt} = \frac{(k_{cat_{dfb}}(b*(1-d))}{(K_{m_{dfb}}+1-d))} - \frac{(k_{cat_{drc}}*c*d)}{(K_{m_{drc}}+d)}$$

icFF

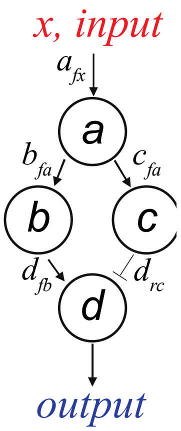


#16

---

Motif #17: Incoherent Feedforward(icFF)

icFF

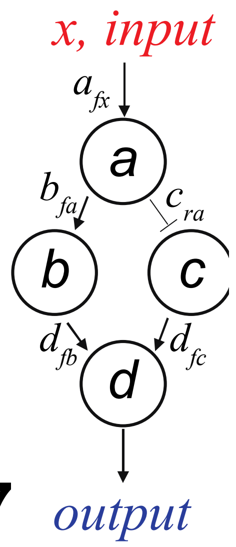


#17

$$\frac{da}{dt} = \frac{(k_{cat_{afx}}\boldsymbol{x}(1-a)}{(K_{m_{afx}}+1-a)} - \frac{(k_{cat_{ar}}(0.5*a))}{(K_{m_{ar}}+a))}$$

$$\frac{db}{dt} = \frac{(k_{cat_{bfa}}(a*(1-b))}{(K_{m_{bfa}}+1-b)} - \frac{(k_{cat_{br}}*0.5*b))}{(K_{m_{br}}+b)}$$

$$\frac{dc}{dt} = \frac{(k_{cat_{cf}}(0.5*(1-c))}{(K_{m_{cf}}+1-c))} - \frac{(k_{cat_{cra}}*a*c))}{(K_{m_{cra}}+c))}$$

$$\frac{dd}{dt} = \frac{(k_{cat_{dfb}}(b*(1-d))}{(K_{m_{dfb}}+1-d))} + \frac{(k_{cat_{dfc}}(c*(1-d))}{(K_{m_{dfc}}+1-d))} - \frac{(k_{cat_{dr}}*0.5*d))}{(K_{m_{dr}}+d))}$$

---

Motif #18: Negative Coherent Feedforward (ncFF)

$$\frac{da}{dt} = \frac{\left(k_{cat_{afx}} \boldsymbol{x} (1-a)\right)}{\left(K_{m_{afx}} + 1 - a\right)} - \frac{\left(k_{cat_{ar}} (0.5 * a)\right)}{\left(K_{m_{ar}} + a\right)}$$

$$\frac{db}{dt} = \frac{\left(k_{cat_{bfa}} \left(a * (1-b)\right)\right)}{\left(K_{m_{bfa}} + 1 - b\right)} - \frac{\left(k_{cat_{br}} * 0.5 * b\right)}{\left(K_{m_{br}} + b\right)}$$

$$\frac{dc}{dt} = \frac{\left(k_{cat_{cf}} \left(0.5 * (1-c)\right)\right)}{\left(K_{m_{cf}} + 1 - c\right)} - \frac{\left(k_{cat_{cra}} * a * c\right)}{\left(K_{m_{cra}} + c\right)}$$

$$\frac{dd}{dt} = \frac{\left(k_{cat_{dfc}} \left(c * (1-d)\right)\right)}{\left(K_{m_{dfc}} + 1 - d\right)} - \frac{\left(k_{cat_{drb}} * b * d\right)}{\left(K_{m_{drb}} + d\right)}$$

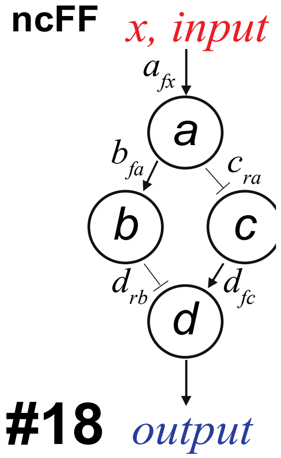


---

Motif #19: Coherent Feedforward (cFF)

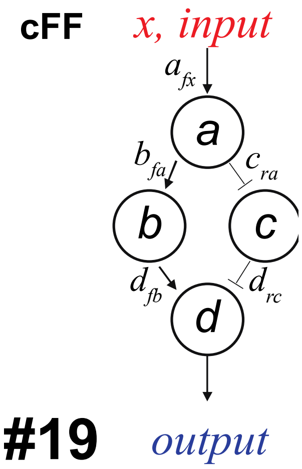


$$\frac{da}{dt} = \frac{\left(k_{cat_{afx}} \boldsymbol{x} (1-a)\right)}{\left(K_{m_{afx}} + (1 - a)\right)} - \frac{\left(k_{cat_{ar}} (0.5 * a)\right)}{\left(K_{m_{ar}} + a\right)}$$

$$\frac{db}{dt} = \frac{\left(k_{cat_{bfa}} \left(a * (1-b)\right)\right)}{\left(K_{m_{bfa}} + 1 - b\right)} - \frac{\left(k_{cat_{br}} * 0.5 * b\right)}{\left(K_{m_{br}} + b\right)}$$

$$\frac{dc}{dt} = \frac{\left(k_{cat_{cf}} \left(0.5 * (1-c)\right)\right)}{\left(K_{m_{cf}} + 1 - c\right)} - \frac{\left(k_{cat_{cra}} * a * c\right)}{\left(K_{m_{cra}} + c\right)}$$

$$\frac{dd}{dt} = \frac{\left(k_{cat_{dfb}} \left(b * (1-d)\right)\right)}{\left(K_{m_{dfb}} + 1 - d\right)} - \frac{\left(k_{cat_{drc}} * c * d\right)}{\left(K_{m_{drc}} + d\right)}$$

---

Motif #20: Negative Coherent Feedforward (ncFF)

$$\frac{da}{dt} = \frac{(k_{cat_{afx}}\boldsymbol{x}(1-a))}{(K_{m_{afx}}+1-a))} - \frac{(k_{cat_{ar}}(0.5*a))}{(K_{m_{ar}}+a)}$$

$$\frac{db}{dt} = \frac{(k_{cat_{bfa}}(a*(1-b))}{(K_{m_{bfa}}+1-b))} - \frac{(k_{cat_{br}}*0.5*b))}{(K_{m_{br}}+b))}$$

$$\frac{dc}{dt} = \frac{(k_{cat_{cfa}}(a*(1-c))}{(K_{m_{cfa}}+1-c))} - \frac{(k_{cat_{cr}}*0.5*c))}{(K_{m_{cr}}+c))}$$

$$\frac{dd}{dt} = \frac{(k_{cat_{df}}(0.5*(1-d))}{(K_{m_{df}}+(1-d))} - \frac{(k_{cat_{drb}}*b*d)}{(K_{m_{drb}}+d)} - \frac{(k_{cat_{drc}}*c*d))}{(K_{m_{drc}}+d)}$$

**ncFF**

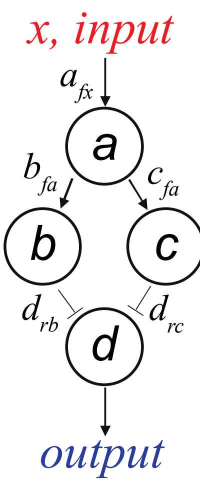


**#20**
