## Supplementary material for "Biochemical network motifs can transduce and process oscillatory information": S3 Appendix - Output Characteristic Equations

**S3 Appendix.** Equations used to calculate output characteristics for each simulation. Three criteria were used to select sustained and transient responders: absolute DC shift, dB suppression, and maximum response.

Absolute DC shift:

The average output value for the last 3 periods of oscillations was calculated using the period of the input signal. This average value was subtracted from the initial output value. The difference was the absolute DC shift.

Output( $i_f$ ) is the average value for last 3 periods of oscillations, $i_f$ is the range of values over which output( $i_f$) is calculated, and $i_o$ is the initial value of the output.

Maximum Response

First, the absolute value of the difference in amplitude of the output and the system's initial amplitude was calculated at each timepoint (t>0). The maximum response was defined as the largest difference in magnitude.

dB suppression:

The fast Fourier transform (fft) of the output signal with time vector t≥ 0 was taken for each individual simulation. The Fourier transform was then divided by the length of the time-vector where t≥0, followed by a final division by 0.2.

The equation used to calculate dB suppression in MATLAB code form can be expressed as: **db suppression = abs(fft(output(t>=0,(k-1)*N+ii),NFFT,1)/L/0.2);**
