## Supplementary material for "Biochemical network motifs can transduce and process oscillatory information": All Supplementary Figures

**S1 Fig.** Graphical representation of the 20 network motifs.

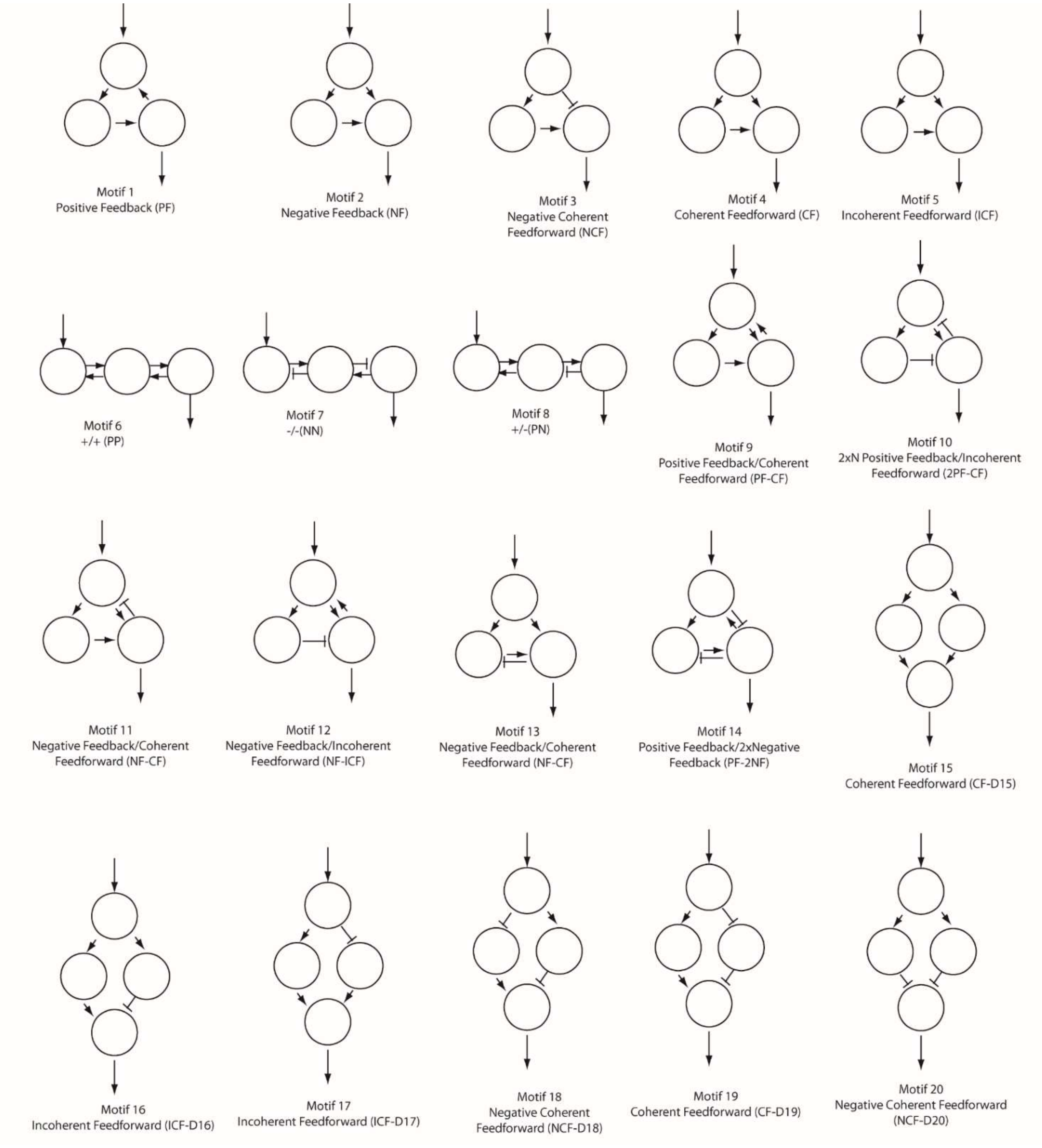

**S4 Fig.** Oscillatory suppression (dB) probability of occurrence for 20 network motifs at all seven frequencies.

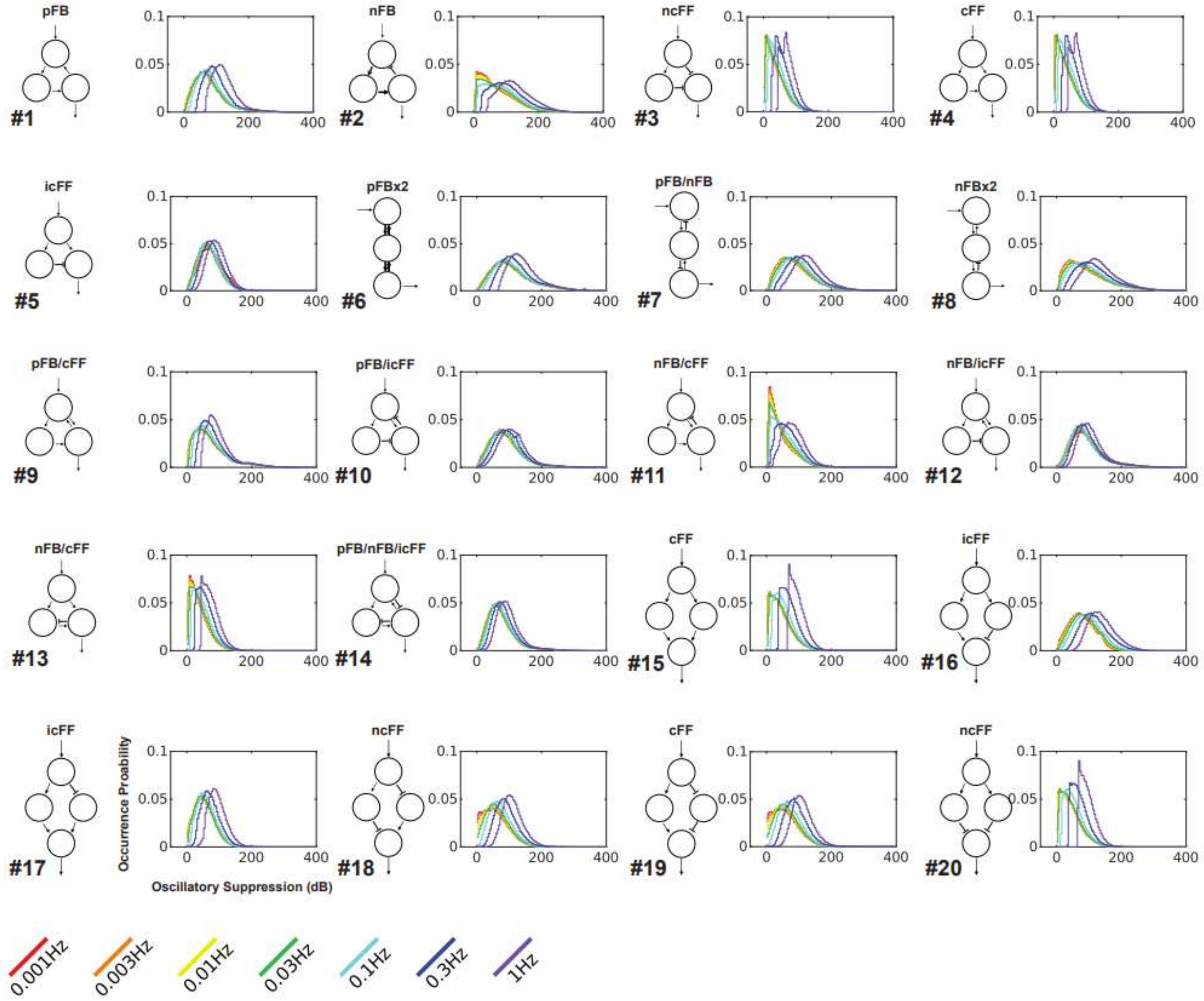


**S5 Fig.** Absolute DC shift probability of occurrence for 20 network motifs at all seven frequencies.

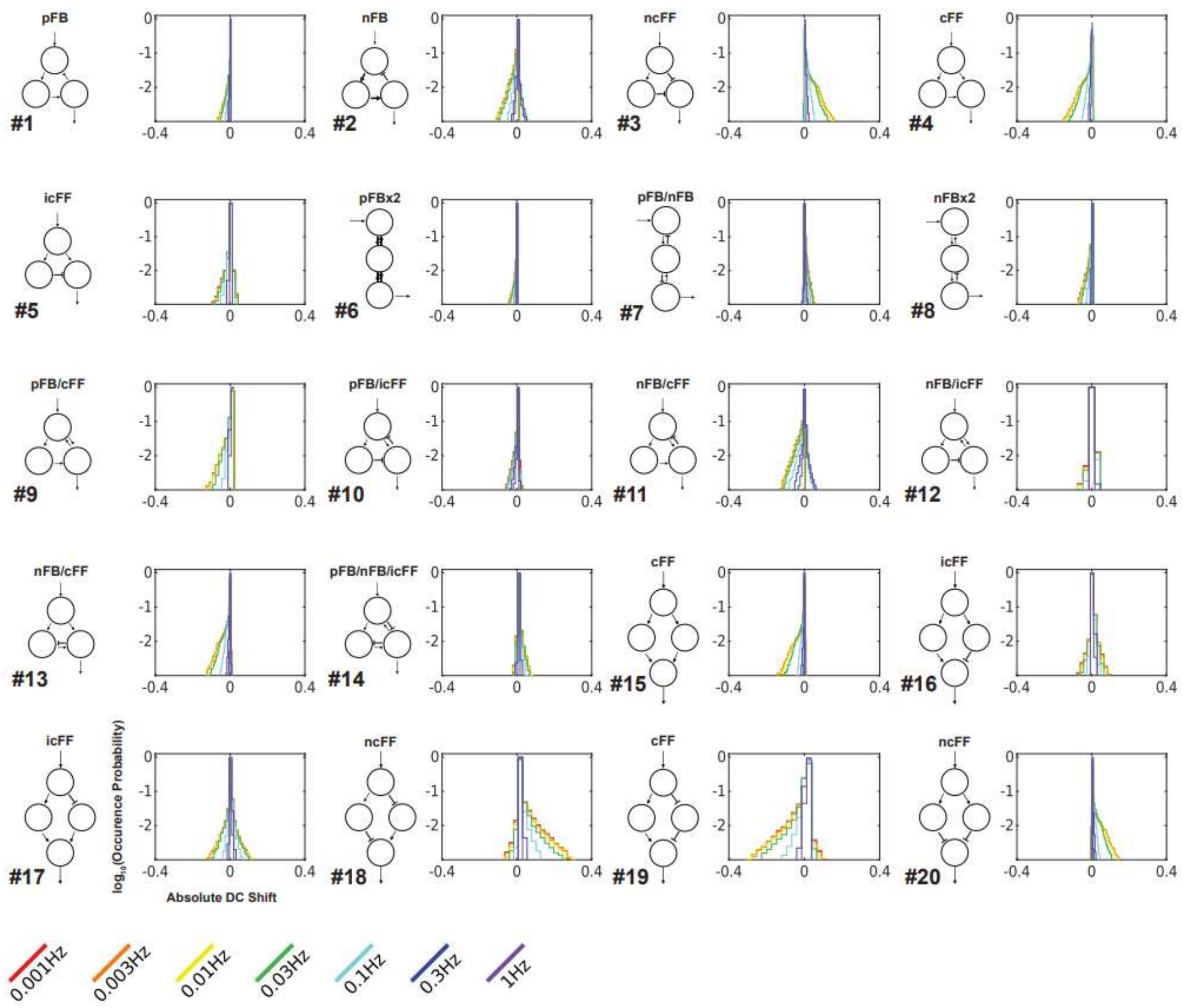


**S6 Fig.** Comparison of enzyme parameters of Motif 19 which produced positive-shifted sustained responses. **(A)** Positive-shifted sustained output responses transduced through Motif 19 at various frequencies. **(B)** Enzyme parameters of Motif 19 which produced each sustained response in (A). **(C)** An equal number of non-sustained response producing parameter sets selected from Motif 19 simulations.

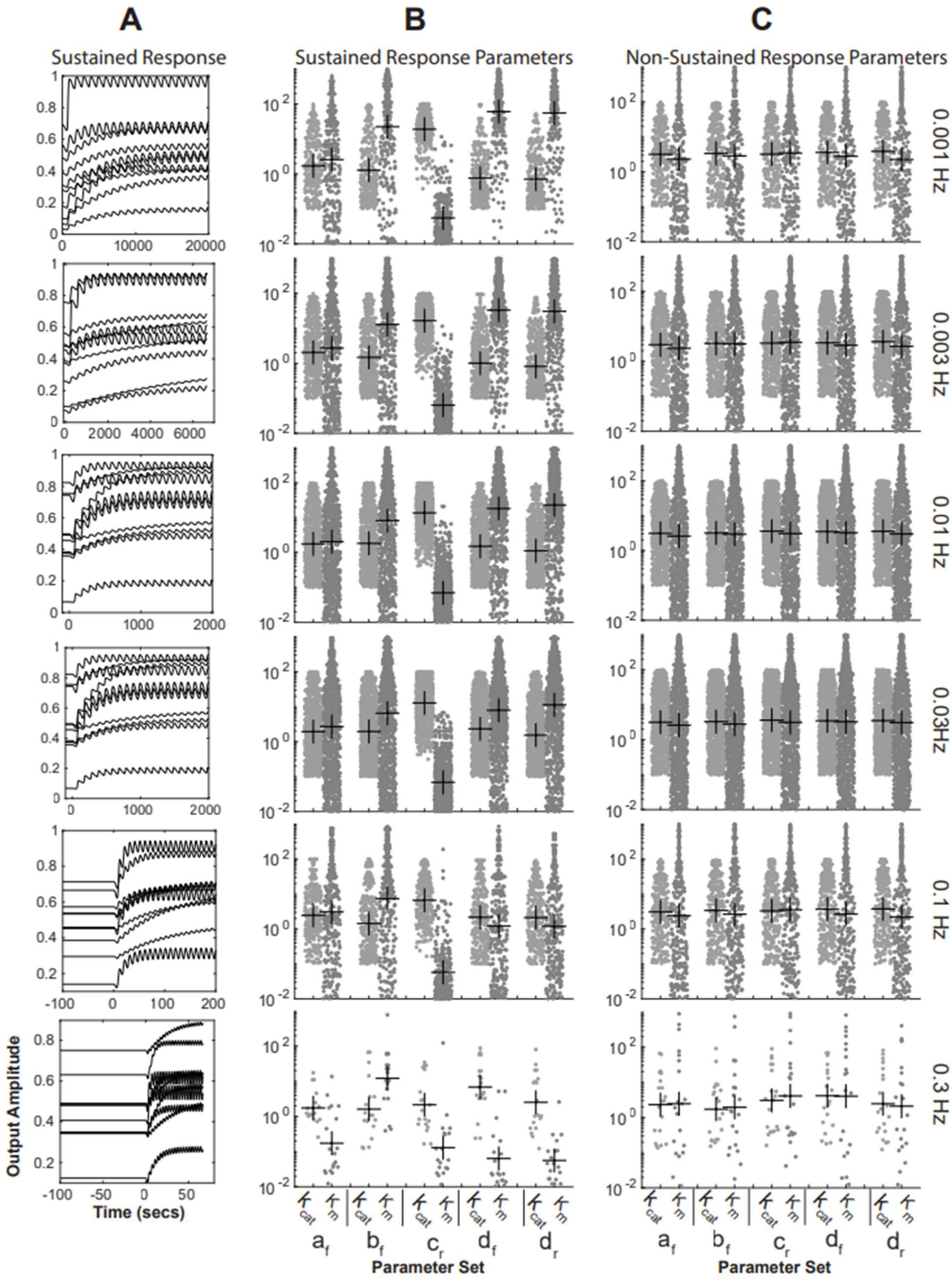

A
B
C
Sustained Response
Sustained Response Parameters
Non-Sustained Response Parameters
0.001 Hz
0.003 Hz
0.01 Hz
0.03Hz
0.1 Hz
0.3 Hz
Output Amplitude
Time (secs)
Parameter Set
Parameter Set

**S7 Fig.** Parameter sensitivity sweep of Motif 19 looking for transient responders. Parameters which produced a transient output in response to a oscillatory input signals are plotted.

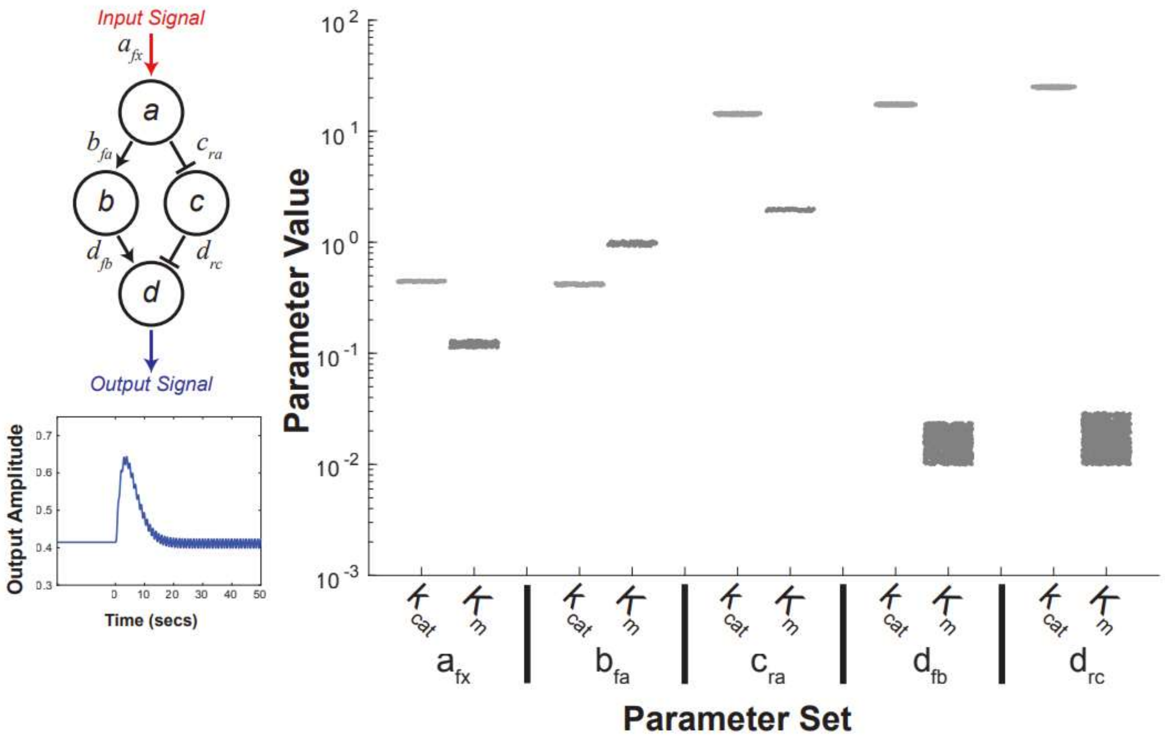
